# Mechanically induced nuclear shuttling of β-catenin requires co-transfer of actin

**DOI:** 10.1101/2021.11.22.469581

**Authors:** Buer Sen, Zhihui Xie, Sean Howard, Maya Styner, Andre J van Wijnen, Gunes Uzer, Janet Rubin

## Abstract

Mesenchymal stem cells (MSC) respond to environmental forces with both cytoskeletal re-structuring and activation of protein chaperones of mechanical information, β-catenin and Yes-Associated Protein 1 (YAP1). To function, MSCs must differentiate between dynamic forces such as cyclic strains of extracellular matrix due to physical activity and static strains due to ECM stiffening. To delineate how MSCs recognize and respond differently to both force types, we compared effects of dynamic (200 cycles x 2%) and static (1 × 2% hold) strain on nuclear translocation of β-catenin and YAP1 at 3h after force application. Dynamic strain induced nuclear accumulation of β-catenin, and increased cytoskeletal actin structure and cell stiffness, but had no effect on nuclear YAP1 levels. Critically, both nuclear actin and nuclear stiffness increased along with dynamic strain-induced β-catenin transport. Augmentation of cytoskeletal structure using either static strain or lysophosphatidic acid (LPA) did not increase nuclear content of β–catenin or actin, but induced robust nuclear increase in YAP1. As actin binds β-catenin, we considered whether β-catenin, which lacks a nuclear localization signal, was dependent on actin to gain entry to the nucleus. Knockdown of cofilin-1 (*Cfl1*) or importin-9 (*Ipo9*), which co-mediate nuclear transfer of G-actin, prevented dynamic strain-mediated nuclear transfer of both β-catenin and actin. In sum, dynamic strain induction of actin re-structuring promotes nuclear transport of G-actin, concurrently supporting nuclear access of β-catenin via mechanisms utilized for actin transport. Thus, dynamic and static strain activate alternative mechanoresponses reflected by differences in the cellular distributions of actin, β-catenin and YAP1.

**Significance statement:** Cells integrate both static and dynamic mechanical signals through the actin cytoskeleton which is attached to the nuclear envelope, affecting nuclear transport of β-catenin and YAP1. Dynamic strain induces nuclear translocation of β-catenin, but not YAP1, while static strain causes nuclear translocation of YAP1, but not β-catenin. Importantly, nuclear transport of actin is induced by dynamic but not static force. Furthermore, nuclear import of β-catenin depends on cofilin/importin-9 dependent actin transport mechanisms. Thus the presence of β-catenin and YAP1 in the nucleus represent specific responses to regulatory mechanical signals.

## Introduction

Mechanical information, both extrinsic and intrinsic to the cell, regulates proliferation, differentiation and function. β-catenin and YAP1 are mechanoresponsive proteins that modulate these processes in mesenchymal stem/stromal cells (MSCs)^1–3^. β-catenin nuclear localization occurs after dynamic strain^1, 4–6^, which concurrently induces actin rearrangement into F-actin cables which arch over the nucleus^7^ and connect to the inner nuclear membrane through actin’s association with the Linker of Nucleoskeleton and Cytoskeleton (LINC) complex^8^. YAP1 nuclear translocation is also modulated by cytoskeletal actin fibers which form in response to forces generated after adherence to stiff substrates^2^, but has not been studied with respect to dynamic strain. Static strain, in turn, has not been shown to effect rapid nuclear β-catenin transfer^9^. Overall, how the cell regulates nuclear import of β-catenin and YAP1 in response to acute application of dynamic or static mechanical force is poorly defined.

Both mechanoresponsive molecules, β-catenin and YAP1, lack nuclear localization signals in their primary structure^10, 11^ and are subject to phosphorylations guiding their disposition. For example, phosphorylation of β-catenin by GSK3β targets it for proteasomal degradation^4^, while phosphorylation of YAP1 by LATS1/2 leads to its exclusion from the nucleus^12^. Mechanical force, through AKT-mediated phosphorylation of GSK3β, inhibits β-catenin turnover and enhances nuclear import^4, 13^. However, these phosphorylation events do not fully explain nuclear transfer engendered by mechanical force. YAP1 nuclear import through nuclear pores is enhanced by actin cables which develop after static strain^14^. In turn, when F-actin connections to the nuclear actin cap decrease due to mechanical unloading (e.g., simulated microgravity), nuclear YAP1 levels decrease^15, 16^. The actin cytoskeleton that forms after dynamic strain also connects to the nucleus via LINC, where β-catenin positions itself for entry^17^. While this might suggest that entry of β-catenin into the nucleus is also consequent to cytoskeletal actin structure, nuclear localization of YAP1 and β-catenin separate after cytochalasin D induced disruption of actin polymerization: cytochalasin D leads YAP1 egress, but promotes nuclear accumulation of β-catenin^18^. It thus appears that nuclear translocation of β-catenin in response to dynamic force is achieved through a fundamentally distinct mechanism than that controlling the entry of YAP1.

Biomechanical forces are represented by a continuum. Tissues respond to these forces by integrating distinct types of biophysical input that differ in frequency (periodicity) and amplitude (magnitude) in a manner that depends on biological and cellular contexts^19^. YAP1 related mechanoresponses have been largely examined as a late response to changes in the static modulus of the substrate on which cells adhere^2, 14^, while β-catenin has been studied as an acute response to dynamic strain regimens^20^. The responses of YAP1 and β-catenin together were examined in a canine kidney cell line in which altered modulus was replaced with a single biaxial stretch of high magnitude^21^. This type of force application resulted in the translocation of both mechanosensory proteins to the nucleus from prior sites in the cytoplasm (YAP1) or cell-cell contacts (β-catenin). In these epithelial cells, YAP1 moved rapidly into the nucleus, but β-catenin nuclear entry required at least 6 hours. The activation of both proteins was associated with entry into the cell cycle, similar to the cell cycle response of MSC to β-catenin nuclear entry^22^. The latter findings suggest functional specialization or perhaps redundancy in how mechanoresponders might regulate cell proliferation.

Because both β-catenin^23, 24^ and YAP1^25^ lack classic nuclear localization signals (NLS) we postulated that assessment of their responses to distinct mechanical forces would offer clues to mechanisms of nuclear transfer. The results presented here show that both β-catenin and YAP1 respond to acute application of strain but exhibit differences with respect to periodicity of the mechanical signal. Furthermore, nuclear actin content increased after dynamic strain, but not static, strain. Importantly, we found that nuclear entry of β-catenin depends on concurrent nuclear entry of actin. In sum, mechanically induced nuclear entry of β-catenin differs from that of YAP1 as it is independent of cytoplasmic actin structure, depending instead on mechanisms of nuclear actin transport.

## Methods

### Materials

Fetal bovine serum was from Atlanta Biologicals (Atlanta, GA). Culture media, trypsin-EDTA reagent, antibiotics from Sigma-Aldrich (St. Louis, MO). Other commercial reagents include 1-Oleoyl Lysophosphatidic Acid (LPA; cat# 62215) from Caymen Chemical (Ann Arbor, MI), 6-well BioFlex Collagen-I coated plates (cat# BF-3001C) from Flexcell International (Burlington, NC), Ibidi USA μ-Slide chamber (cat#: NC0515977) from Fisher (Hampton, NH).

### Cells and culture conditions

Mouse marrow-derived MSCs were harvested from murine bone marrow using a published protocol^26^. Cells were maintained in minimal essential medium (MEM) containing 10% FBS, 100 μg/ml penicillin/streptomycin. For immunofluorescence staining, MSCs were plated at a density of 2,000 (β-catenin) or 10,000 (YAP1) cells per square centimeter and cultured for 1 day before application of treatments. For YAP1 staining, the cells were cultured in adipogenic medium (0.1 μM dexamethasone, 50 μM indomethacin, 5 μg/ml insulin) overnight before mechanical loading or reagent treatment.

### RNA interference

Cells were transfected with siRNA (50 nM) in serum-free OptiMEM overnight before replacing the medium and adding reagents for cell treatment. RNA interference studies were performed with siRNAs for Ipo9: 5’-CCCAGCUCUUCAACCUGCUUAUGGA and control (nucleotide change within same sequence) 5’-CCCTCTCCTAACCGTTCATTGAGGA; Cfn1: 5’-AAACTAGGTGGCAGCGCCGTCATTT and the control 5’-TCATTTCCCTGGAGGGCAAGCCTTT.

### Application of mechanical forces

Uniform biaxial strain was applied to MSCs plated at 10,000 cells per well on 6-well Bioflex Collagen-I coated plates using the Flexcell FX6000T system (Flexcell International, Hillsborough, NC). The mechanical regimen consisted of 2% dynamic strain at 10 cycles per minute for 20 minutes (200 cycles) or 2% static strain for 3h continuously (1 cycle). For adipogenic differentiation, primary bone marrow derived murine MSCs were seeded on BioFlex culture plates at 10,000 cells per well 2 days prior to addition of adipogenic media (5 μg/ml insulin, 0.1 μM Dexamethasone, and 50 μM Indomethacin). Cells were incubated overnight and then subjected to either dynamic strain (2% for 200 cycles) or static strains (2% continuously for 1 cycle) using a Flexcell FX5000 Tension device.

### Measurement of cell and nuclear modulus

For cell modulus measurement, the silicon base of the culture plate was excised with a scalpel after Flexcell strain treatment. Specimens were carefully transferred to the lid of a 35mm culture plate for analysis by AFM. Nuclei were isolated from strain plates using a NEPER nuclear isolation kit (Thermo Scientific) according to manufacturer protocols. Force-displacement curves for both intact MSC nuclei and isolated nuclei were acquired using a Bruker Dimension FastScan Bio AFM. MLCT-SPH-5um -DC-A probes were used to decrease variables in calculating moduli. These probes are produced with a consistent 5 um silicon nitride tip and factory calibrated, using laser doppler vibrometry, spring constants. To fully calibrate the probes, the deflection sensitivity was measured immediately prior to each experiment by ramping on sapphire and averaging five measurements. MSCs and nuclei were located using the AFM’s optical microscope and engaged on using a minimal force set point (1-3 nN) to ensure contact while minimizing applied force and resultant deformation prior to testing. Ramps were performed over the approximate center of each nucleus for all samples. After engaging on a selected nucleus to ensure probe/nucleus contact as described above, force curve ramping was performed at a rate of 2 μm/sec over 2 μm total travel (1 μm approach, 1 μm retract). 5 replicate force-displacement curves with an indentation depth of ^~^500nm were acquired and saved for each nucleus tested, with at least 3 seconds of rest between ramps.

Measured force-displacement curves were analyzed assuming Hertzian (spherical contact) mechanics^27^. AtomicJ, an open source program for analyzing AFM data was used to automatically fit the force-displacement curves and calculate the modulus^28^. Poisson’s ratio 0.4 was used as a best estimate for the effective Poisson’s ratio of the cells. The modulus of each individual sample was reported as the average of at least five consecutive force curve measurements. Group averages were then obtained from the average values for the individual samples.

### Immunofluorescence microscopy

Cells were fixed with 4% paraformaldehyde for 10 minutes, permeabilized in 0.1% Triton-X 100 for 5 minutes, and blocked with 5% donkey serum for 30 minutes. Three consecutive washes for 10-minutes with phosphate-buffered saline (PBS) were performed between each step. Silicone membranes were cut from plates and transferred to six-well plate surface. Incubation with primary antibodies at 37°C for 1h included goat β-catenin antibody (cat# AB0095) from OriGene Biotechnologies (Rockville, MD) and rabbit YAP1 antibody (cat# 14074) from Cell Signaling (Danvers, MA). Visualization of primary antibodies was performed with Cy5 conjugated donkey anti-goat (for β-catenin; cat# 705-175-147) and Rhodamine Red-conjugated anti-rabbit (for YAP1; cat# 711-295-152). These secondary antibodies were obtained from Jackson Immuno Research (West Grove, PA). Actin stress fibers were examined using Alexa Fluor 488-conjugated phalloidin (cat# A12379; Lifetechnologies). After three consecutive 10-minute washes with phosphate-buffered saline (PBS), membranes were sealed with mounting medium on glass or the cells in chamber slides were covered with PBS. For 3D images, cells were examined using a model LSM 880 confocal microscope (Zeiss, Thornwood, NY).

### Real Time RT-PCR

Total RNA was isolated with the RNeasy plus mini kit (Qiagen; Germantown, MD). Reverse transcription of 1 μg of RNA in a total volume of 20 μl was performed with iScript cDNA Synthesis Kit (Bio-Rad) prior to real time PCR (iCycler; BioRad, Hercules, CA). 25-μl amplification reactions contained primers (0.5 μM), dNTPs (0.2 mM each), 0.03 units Taq polymerase, and SYBR-green (Molecular Probes, Eugene, OR) at 1:150,000. Aliquots of cDNA were diluted 5 to 5000-fold to generate relative standard curves to which sample cDNA was compared. *Cfl1* forward primer: 5’-cgcaagtcttcaacaccaga-3’ and reverse primer: 5-ttgtctggcagcatcttgac-3’. *Ipo9* forward primer: 5’-tggattcggatggagaagtc-3’ and reverse primer: 5’-agtaaatgagctcgggcaga-3’. Standards and samples were run in triplicate. PCR products were normalized to 18 S amplicons in the RT sample, and standardized on a dilution curve from RT sample.

### Nuclear and cytoplasmic protein fractionation

Cells were washed with 1x PBS, cell pellets resuspended in 0.33 M sucrose, 10mM Hepes, pH 7.4, 1mM MgCl_2_, 0.1% Triton X-100 (pellet *versus* buffer volume, 1:5) and placed on ice for 15 min. After 3,000 rpm for 5 min, the supernatant was collected (cytoplasmic fraction). Pellets were resuspended in 0.45 M NaCl and 10mM Hepes, pH 7.4, and placed on ice for 15 min. After centrifugation at 12,000 rpm for 5 min, the nuclear fraction supernatant was collected.

### Immuno-precipitation (IP) analysis

MSCs were transfected with recombinant plasmid vectors that express either a chimeric actin protein fused to fluorescent YFP and a nuclear localization signal (NLS peptide) or the YFP-NLS control protein. Cells were separated into cytoplasmic and nuclear compartments fractions that were incubated with GFP antibody (cat# 2555, Cell Signaling) which recognizes the YFP protein. Immuno-complexes were bound to Protein A/G beads and recovered by low-speed micro centrifugation at 3000rpm (approximately 1000X g) for 30 seconds at 4C. After final wash, pellets were resuspended in PBS for SDS-PAGE analysis.

### Immunoblot analysis

Fractionated proteins were loaded onto a 7–10% poly-acrylamide gel for chromatography and transferred to polyvinylidene difluoride membrane. After blocking, primary antibody was applied overnight at 4 °C including antibodies against β-catenin (cat# AB0095-200, OriGene), β-Actin (cat# 4967, Cell Signaling), GFP (cat#2555, Cell Signaling), PARP1 (cat# sc-365315, Santa Cruz Biotechnology, Dallas, TX,) or LDHA (cat# 2012, Cell Signaling). Secondary antibody conjugated with horseradish peroxidase was detected with ECL plus chemiluminescence kit (Amersham Biosciences/GE Healthcare, Piscataway NJ). The images were acquired with an HP Scanjet and densitometry determined using NIH ImageJ, 1.37v.

### Statistical analysis

The results are expressed as means ± standard error of the mean (SEM). For comparisons, one-way analysis of variance or *t* test (GraphPad Prism). Experiments were replicated at least twice to assure reproducibility. Densitometry data, where given, were compiled from at least three separate experiments. EFM experiments were repeated x 3. P-values of < 0.05 were considered significant.

## Results

### Dynamic and static strains both increase cell stiffness, yet have different effects on mechanoresponders β-catenin and YAP1

YAP1’s mechanoresponse has been studied after plating on substrates of high stiffness (>40kPa), as noted in bone marrow MSC, mammary epithelial and HeLa cervical carcinoma cells^2^. YAP1 also responds to a single stretch of 15% within a 2 hour time frame in MDCK kidney epithelial cells^21^. In contrast, application of physiologically relevant dynamic strain (200 cycles x 2%) to bone marrow MSCs induces rapid nuclear transfer of β-catenin^4, 29^. Here we compared the specific responses of β-catenin and YAP1 to physiological levels of dynamic and static strains in mouse MSCs. Application of 200 cycles of 2% strain is referred to here as “dynamic” strain, while a similar magnitude strain, applied and held at 2%, is referred to as “static” strain. Phalloidin staining showed an increase in F-actin in both dynamic and static conditions at 3 hours (**Fig 1A**). In order to measure changes in cell stiffness, MSCs treated with either dynamic or static strain along with non-strain controls were tested under an atomic force microscope to measure live cell modulus as we have previously reported^30^. On average, application of dynamic strain increased the AFM-measured cell modulus by 22% compared to non-strained controls and 11% compared to static strain, while static strain resulted in a 13% increase compared to non-strained controls (**Fig 1B**).

**Fig 1.**
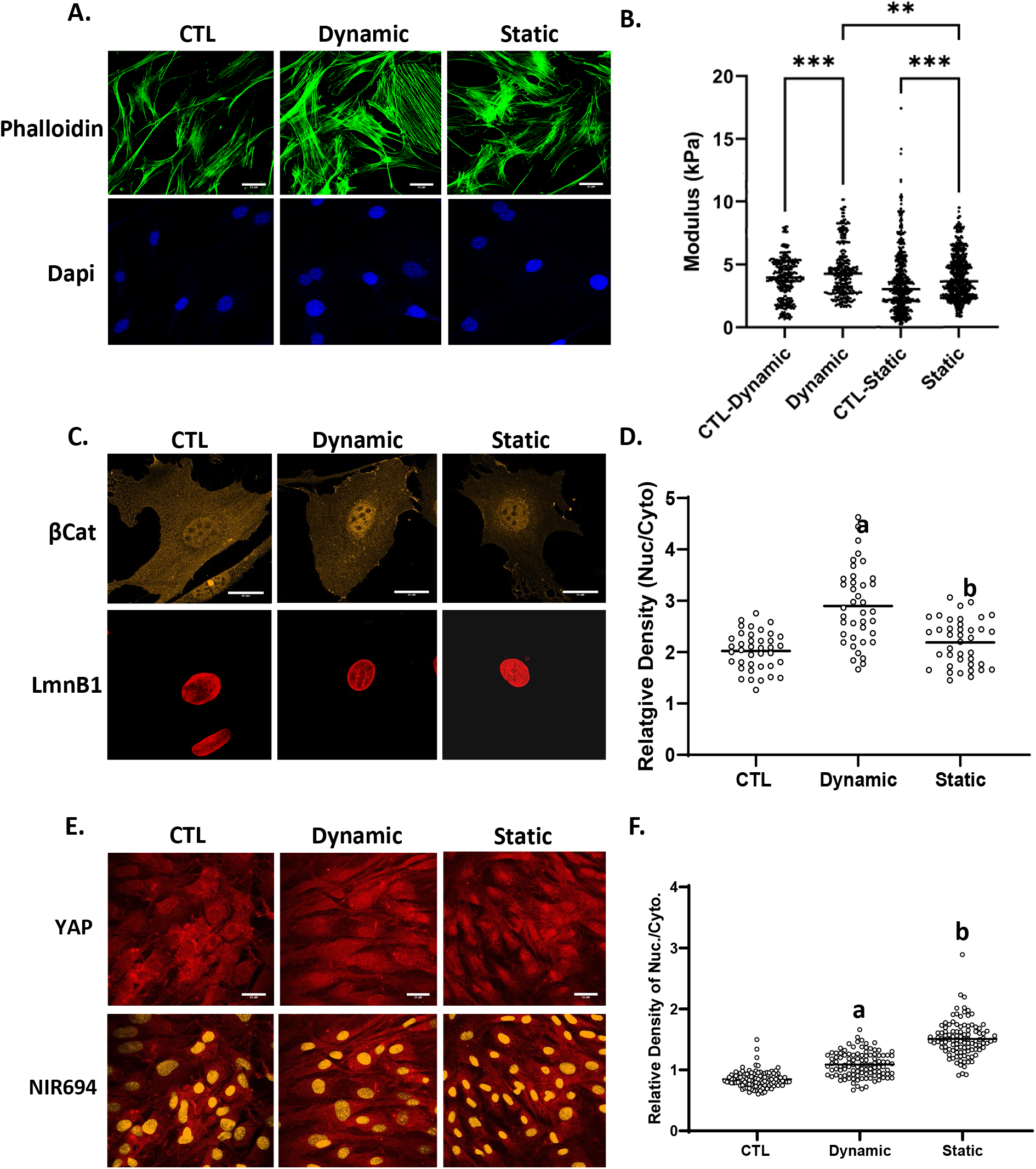
Dynamic and static strains both increase cell stiffness yet have different effects on mechanoresponders. MSCs were seeded in MEM at a density of 2,000 or 10,000 cells/cm^2^ for β-catenin or YAP1 experiments respectively. 24 h later, application of 2% dynamic or static strain was followed by staining with phalloidin (green), anti-β-catenin (yellow) or anti-YAP (red), scale bar = 25 μm (**A, D** and **E**). Nuclear intensity of β-catenin and YAP1 was analyzed by ImageJ program and statistical significance indicated by letter a and b, both of them p < 0.0001 and a ≠ b (**D** and **F**). **(b)** Application of dynamic strain increased the AFM-measured cell modulus by 22% (p<0.001, N=350/grp) compared to non-strained control and while static strain resulted in a 13% increase compared to non-strained control (p<0.0001, N=800/grp). Cell modulus of MSCs treated with dynamic strain were, on average, 11% larger compared to static strain groups (p<0.05).

At 3h after application of dynamic strain, increases in nuclear β-catenin across the cell population were significant. The static condition induced much less, but still significant β-catenin nuclear transfer (**Fig 1C**). Analysis of β-catenin immunofluorescence (IF) signal over multiple nuclei and similar areas of cytoplasm, revealed the magnitude of this effect (143 ± 5.9% vs 108 ± 3.6% for dynamic and static strain respectively, compared to control, **Fig 1D**).

To assess YAP1 in MSCs, cell confluence was increased to ensure that YAP1 could be clearly detected in the cytoplasm (**Fig 1E**). Both static and dynamic strain induced a similar increase in the F-actin cytoskeleton. However, while dynamic strain had only marginal effects on YAP1 nuclear accumulation by 3 hours, static strain robustly induced nuclear transfer of YAP1 (128 ± 2.2% vs 178 ± 3.3% for dynamic and static respectively, compared to control, **Fig 1E-F**). Hence, dynamic and static strain had qualitatively different effects on the mechanoresponses of β-catenin and YAP1.

### Nuclear actin increases after dynamic strain

Increased nuclear transfer of YAP1 associated with plating cells on substrates of high modulus is thought to require release of YAP1 from actin monomers^31^. Accordingly, depolymerization of cytoplasmic F-actin in MSCs by cytochalasin D results in YAP1 exit from the nucleus. In direct contrast, β-catenin enters the nucleus after F-actin depolymerization^18^. The cytochalasin D effect on either mechanoresponder is complicated by mass transport of depolymerized actin monomers from the cytosol into the nucleus; increased levels of nuclear actin, which represents a substrate for nuclear members of the actin toolbox, can support re-polymerization and perhaps modulate nuclear modulus^32^. For this reason, we investigated whether differences in actin response to dynamic and static strains could affect β-catenin or YAP1 nuclear entry.

To localize cellular actin after dynamic and static strain, both of which induce F-actin polymerization in the cytoplasm^29, 33, 34^, we examined the subcellular localization of a chimeric actin protein fused to fluorescent YFP and a nuclear localization signal (NLS peptide). As expected, the YFP-NLS peptide lacking actin was entirely confined to the nucleus and was insensitive to either dynamic or static strain (**Fig S1**). Fusion of the actin protein to the NLS-YFP protein module resulted in considerable redistribution of YFP to the cytoplasm irrespective of the NLS signal (**Fig 2A**). This finding firmly established that the NLS-YFP-actin fusion protein was not restricted to the nucleus. These results indicated that the NLS-YFP-labeled actin protein remained subject to regulators of actin transport and that its cellular localization was partially governed by molecular regulatory mechanisms that control actin entry and exit from the nucleus. After dynamic strain, actin accumulated in the nucleus, in stark contrast to the static condition where the YFP-actin signal did not increase (**Fig 2A**). Both strain conditions consistently induced cytoplasmic actin polymerization, as evidenced by actin cables containing the YFP-actin-NLS molecule. Western blot assessment of YFP-actin showed nuclear accumulation after dynamic strain, but not after static strain (**Fig 2B-C**).

**Fig 2.**
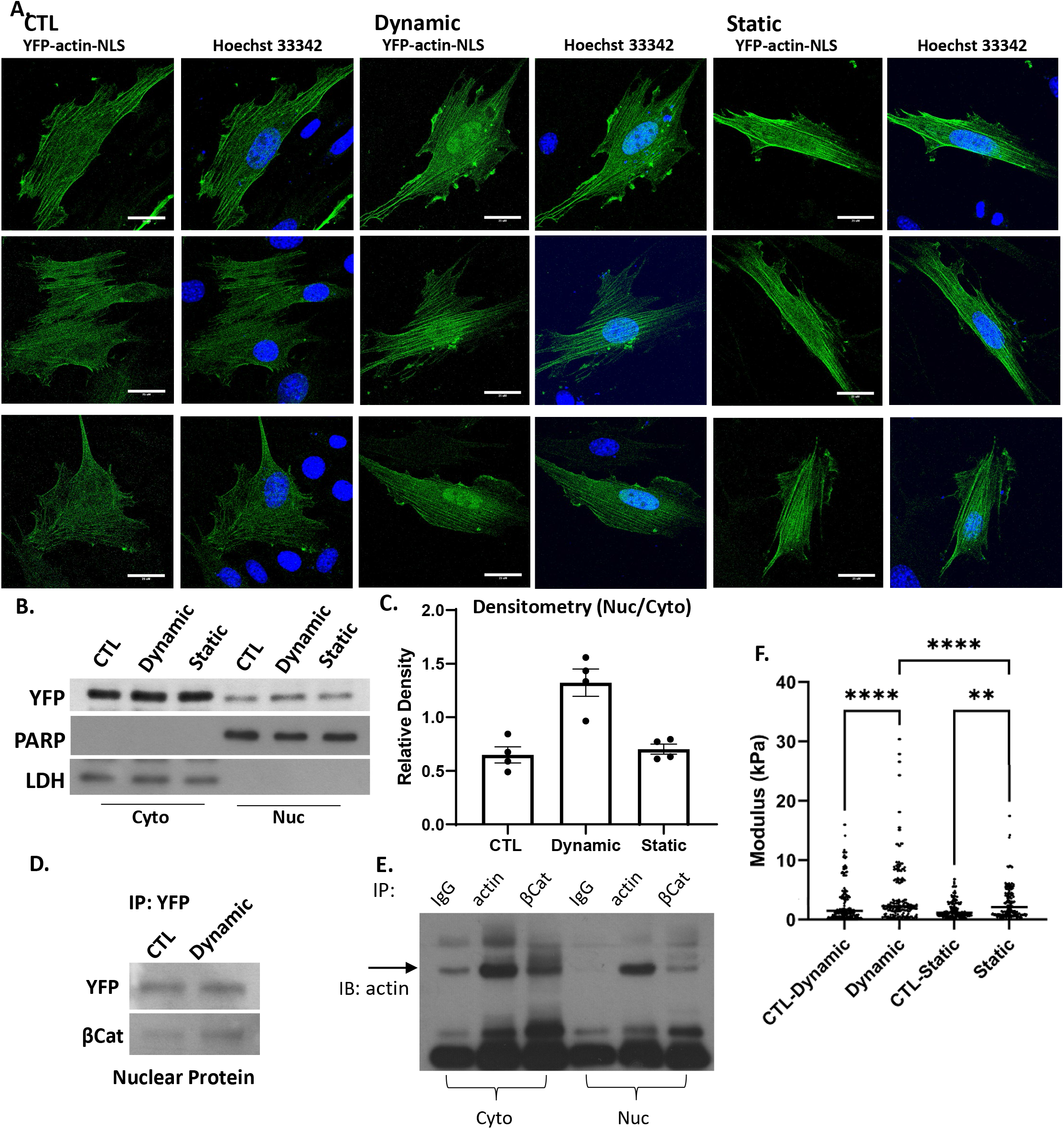
Nuclear actin increases after dynamic strain. Nuclear YFP-NLS-βActin in MSC increases after dynamic, but not static strain (3 examples of each condition shown, scale bar = 25 μm (**A**). Western blot of cytoplasmic and nuclear lysates (**B**) was quantified by densitometry n=4, p < 0.005 (**C**). IP nuclear protein using anti-GFP and IB YFP and β-catenin shows association between the 2 molecules (**D**). Cell cytoplasmic and nuclear lysates were IP for IgG, actin or βCat and immunoblotted for actin (**E**). AFM-measured modulus of isolated nuclei showed that dynamic strain increased the nuclear modulus by 2-fold (p<0.001) compared to control and 1.3-fold compared to static strain groups (p<0.001). Static strain increased nuclear modulus by 1.5-fold compared to control (p<0.05). (**F**).

Actin is known to bind β-catenin/E-cadherin complexes^35^ and when artificially increased in the nucleus, F-actin associates with β-catenin, enhancing its binding to cell cycle regulator genes (i.e., c-myc, cyclin D, OCT4)^36^. After dynamic strain, we measured an increase in nuclear YFP-actin, as well as in the β-catenin associated with the epitope used for immunoprecipitation (**Fig 2D**). As predicted, IP for β-actin pulled down β-catenin in MSCs (**Fig 2E**). Shown in **Fig 2F**, both strain regimens increased the nuclear modulus: dynamic strain increased nuclear modulus 2-fold (p < 0.001) compared to control, and 1.3-fold (P < 0.001) compared to static strain. Static strain increased nuclear modulus by 1.5 fold (p < 0.01) compared to its own control.

Repeated episodes of mechanical force application may sustain and increase effects observed for a single period of mechanical stimuli. As such, we applied a second bout of dynamic strain delivered 3 h after the first bout (**Fig 3A**), a time when augmentation of the F-actin cytoskeleton has plateaued^7^. Micrographic analysis revealed that this force re-application regimen further enhanced β-catenin localization in the nucleus (**Fig 3B,C**), consistent with previous observations^29^. The increased level of nuclear β-catenin was clearly reflected by increased immunofluorescence after the second strain bout (**Fig S1**, 3^rd^ column). Quantitation revealed a significant 30-40% increase in YFP-actin after the first stimulus (136 ± 2.7% compared to control at 100%), and 65-75% after the second episode of strain (170 ± 3.5% compared to control). Immunoprecipitation of the YFP-actin-NLS fusion protein using a GFP antibody revealed that YFP-actin forms stable protein/protein complexes with β-catenin in the nucleus and the level of these complexes increased further after the second bout (**Fig 3D**). In contrast, immunoprecipitation of cytoplasmic YFP-actin did not result in visible β-catenin, perhaps due to cytoplasmic sequestration of β-catenin. These data indicate that β-catenin entry into the nucleus is associated with force-induced actin translocation.

**Fig 3.**
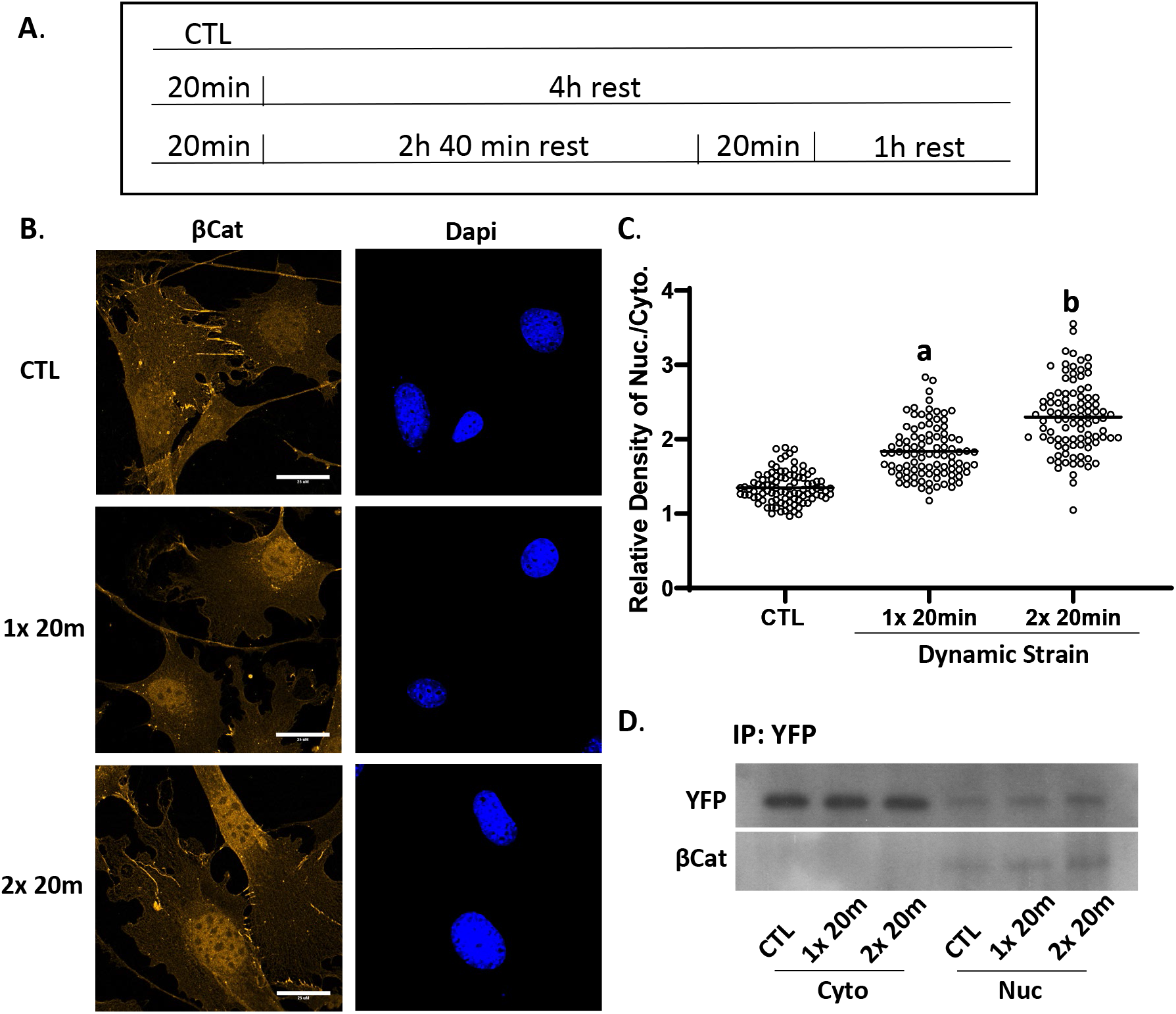
Nuclear β-catenin and actin rise further after a second dynamic strain bout. Timed application of strain for signal and double bouts (**A**). MSC were stained with anti-β-catenin (yellow) or DAPI (blue) showing increased β-catenin signal after the second strain bout, scale bar = 25 μm (**B**). β-catenin was quantified using ImageJ; statistical significance indicated by letter a and b, both = p < 0.0001 and a ≠ b (**C**). Nuclear and cytoplasmic fractions were pulled down with GFP Ab, and Western as shown for YFP and β-catenin, both of which increase after strain application (**D**).

### β-catenin nuclear entry is dependent on active actin transport into the nucleus

Actin transport into the nucleus depends on the co-transport with cofilin-1 and importin-9^18, 37^. When *Cfn1* was depleted by siRNA knock-down, nuclear transport of β-catenin transport after dynamic strain was abolished, while still increased by strain in MSCs treated with the scrambled RNA control (192 ± 6.5% of control; **Fig 4A-C**). Static strain increased nuclear β-catenin to 138 ± 5.5% compared to the unstrained condition in MSCs treated with scrambled RNA. Again, this smaller increase in nuclear β-catenin levels was not observed after *Cfn1* knockdown. There were no effects of *Cfn1* knockdown on static strain levels of nuclear β-catenin, which remained unchanged in both transfection conditions. We next knocked down *Ipo9*, which participates in the nuclear transport of many cargoes including histones^38, 39^ while also acting an obligate co-factor of cofilin-1 in actin transfer^40^ (**Fig 4D-F**). The accumulation of nuclear β-catenin due to dynamic strain evident in cells transfected with scrambled siRNA (158 ±4% compared to control) was completely abolished after *Ipo9* knock down. There was no effect of *Ipo9* knock down on cytosolic β-catenin in the static condition. As anticipated, nuclear transport of YFP-actin-NLS which occurred after dynamic but not static strain, was abolished when *Cfn1* was knocked down (**Fig S2**). These results indicate that mechanically induced nuclear transport of β-catenin is mediated by molecular mechanisms that support nuclear entry of actin.

**Fig 4.**
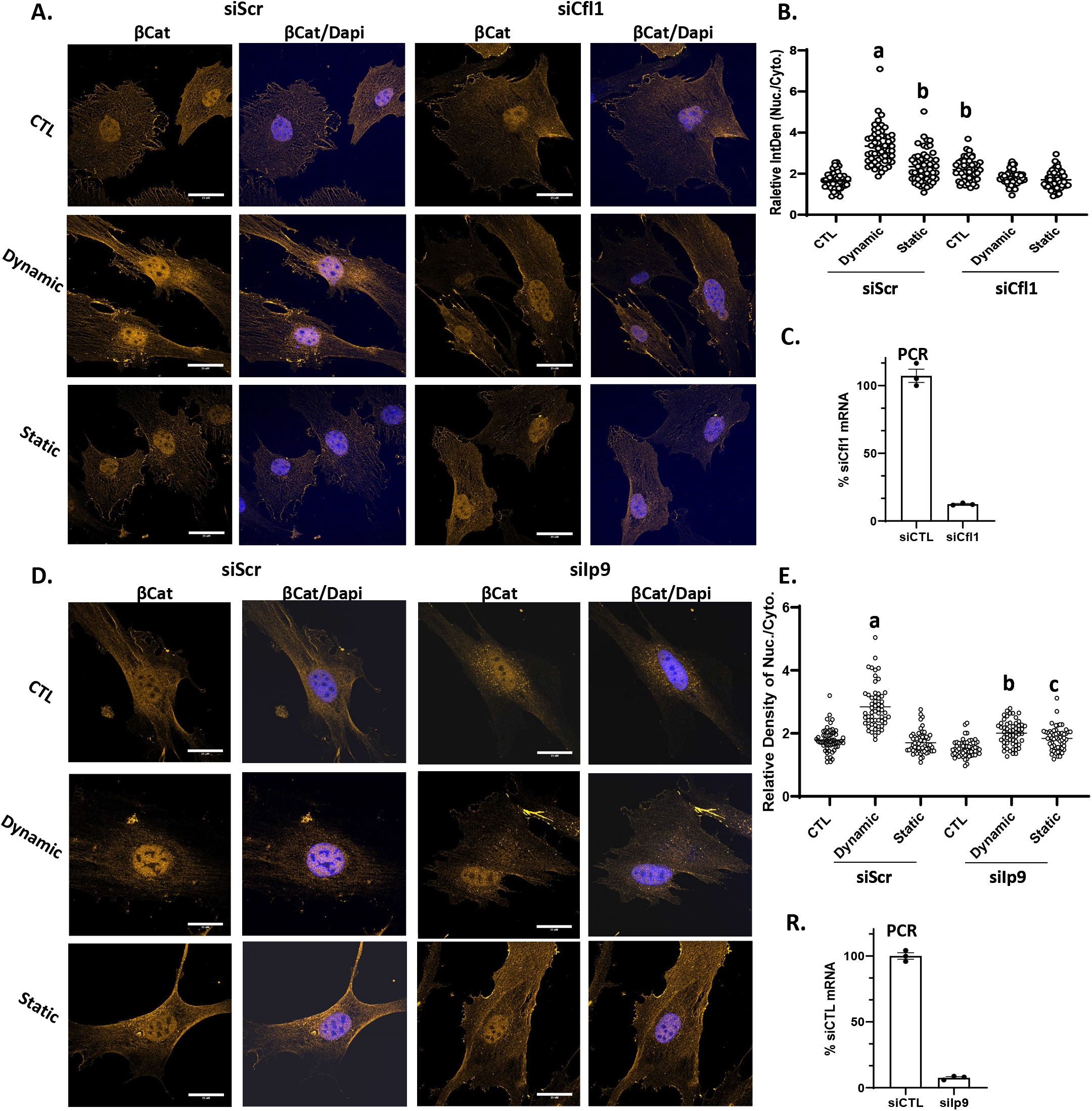
β-catenin nuclear entry is dependent on active actin transport. Cofilin-1 or importin-9 were knocked down in MSC prior to treatment with dynamic or static strain. 3 h later, cells were stained with anti-β-catenin (yellow) and DAPI (blue) (**A** and **D**), scale bar = 25 μm. Nuclear intensity of β-catenin was analyzed by ImageJ program and statistical significance indicated by letter a, b or c. i). a ≠ b, p < 0.0001; ii). b ≠ c; p < 0.05 (**B** and **E**). Real-time PCR indicated knock-down of *Cfl1* or *Ipo9* (**C** and **F**).

### YAP1, but not β-catenin, accumulates in the nucleus after LPA-induced cytoplasmic actin polymerization

Development of F-actin polymers in the cytoplasm due to substrate stiffness or spreading is associated with nuclear transfer of YAP1^2, 31^. Induction of F-actin fibers in the cytoskeleton by ligand-activation of the G-protein coupled lysophosphatidic acid receptor (LPAR1) also induces YAP1 activation^12, 41^. Such cellular redistribution is consistent with the model that YAP1 is sequestered by G-actin and released upon actin polymerization. Similar to previous observations with bone marrow derived MSCs^16^, establishment of an F-actin cytoskeleton by LPA induction was reflected by phalloidin staining and accompanied by YAP1 transfer to the nucleus (193 ± 3.4% compared to control, **Fig 5A,B**). In contrast, LPA ligand activation of LPAR1 and F-actin formation failed to induce relocation of β-catenin from its cytoplasmic and membrane sites into the nucleus. These results indicate that generation of a cytoplasmic actin cytoskeleton was not sufficient to promote nuclear transfer of β-catenin. Rather β-catenin gains access to the nucleus in association with mono- and dimeric actin in response to dynamic mechanical forces.

**Fig 5.**
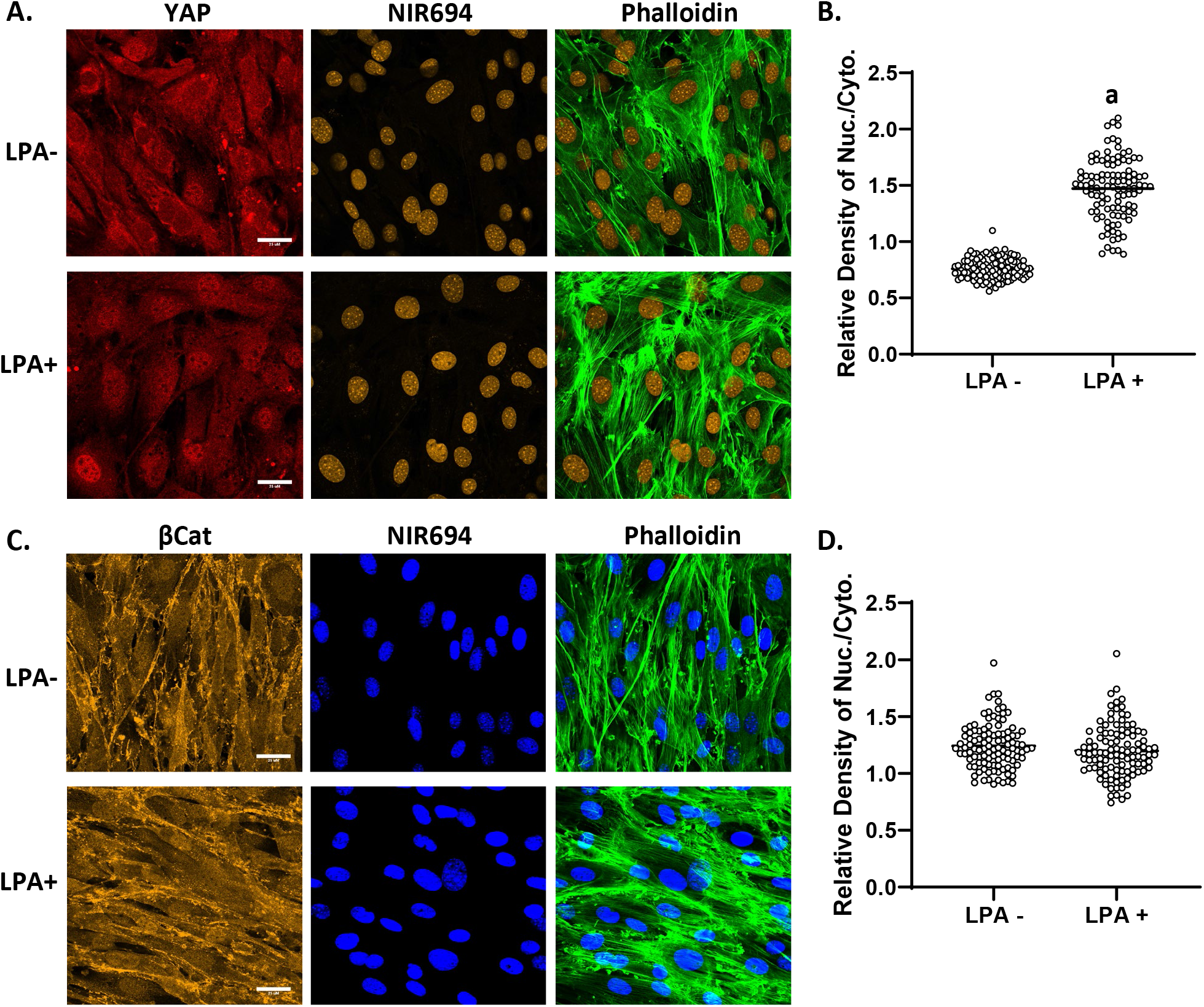
YAP1, but not β-catenin, responds to generation of cytoplasmic F-actin. MSCs at 10,000 cells/cm^2^ were treated with LPA (50 μM) for 24h and stained for YAP1 (red), nucleus (yellow) and F-actin (green) (**A**) or βCat (yellow), nucleus (blue) and F-actin (green), scale bar = 25 μm (**A,C**). Nuclear/cytoplasmic density of YAP1 or β-catenin IF was quantitated by ImageJ, a = p < 0.0001 (**B**, **D**).

## Discussion

β-catenin and YAP1 respond to alterations in the cellular mechanical environment with nuclear translocation and accumulation. The subcellular localization of both proteins is modulated by alterations in the cytoplasmic and nuclear actin state of the cell. Characteristics of the force applied affect these mechanoresponders differently, with dynamic substrate strains driving β-catenin into the nucleus, and static force promoting nuclear transfer of YAP1. Here we show in murine MSCs that nuclear transfer of β-catenin, but not YAP1, requires actin transfer into the nucleus generated by dynamic, but not static strain. Loss of function studies using siRNAs for *Cfn1* and *Ipo9* reveal that β-catenin transfer co-opts the mechanisms responsible for nuclear actin import.

Dynamic strain, through integrin attachments of the cell to substrate, leads to a signaling cascade in which focal adhesion kinase (FAK/PTK2) activates MTOR kinase (as part of the mTORC2 complex), which mediates AKT1 phosphorylation and the subsequent inactivation of GSK3β. The inhibition of GSK3β kinase activity not only protects β-catenin from proteasome destruction, but also enhances β-catenin appearance in the nucleus^4, 13^. The mTORC2 complex and AKT1 also sequentially activate RhoGTPase^42^ to reinforce the presence of signaling molecules at maturing focal adhesions^7^. Such recruitment of cellular proteins to sites of force amplifies actin remodeling^34^ such that reapplication of strain, as shown here, further enhances β-catenin nuclear transfer. Importantly, we show here that dynamic strain generates actin flow into the nucleus. Availability of actin monomers for active transfer to the nucleus may influence the rate of actin remodeling^43^. Furthermore, dynamic strain may influence the iterative activation of RhoGTPase for actin monomer recycling^29^. This process differs from application of static stretch, where the generated cytoplasmic actin structure did not promote significant actin transfer into the nucleus. It is thus possible that nuclear import of actin was responsible for the greater nuclear modulus measured after dynamic, as compared to static, strain. Structural elements such as Lamin A/C^44^ or LINC complex elements Sun 1 &2^45^, as well as chromatin compaction^46^, significantly contribute to nuclei stiffness. Low Intensity Vibration (LIV) which represents another type of dynamic signal, increases isolated nuclear stiffness and leads to chromatin compaction without affecting LaminA/C and Sun proteins levels^30^ suggesting that actin import may also contribute to chromatin organization.

Once inside the nucleus, actin and cofilin have been shown to together form rod structures, especially after mechanical stresses^47^ such as dynamic strain. Formation of nuclear F-actin protects the nucleus from shape deformation, and promotes repair of incipient genetic damage^48^. Moreover, the tendency of actin to organize into secondary filaments can affect cell fate and lineage-progression of MSCs^49^. Thus it is likely that actin, along with β-catenin, might alter heterochromatin architecture and thus modulate transcription. Indeed, we have shown that mass actin transfer due to cytochalasin D treatment induces osteogenic differentiation of MSC^18^, perhaps in part through changes in the expression of Hippo pathway components (e.g., VGLL4)^50^. Furthermore, nuclear β-catenin is capable of preserving the self-renewing multipotent state of MSCs through activation of EZH2^51, 52^, the enzymatic component of the polycomb repressive complex 2 (PRC2) that generates facultative heterochromatin through trimethylation of histone 3 lysine 25 (H3K27me3). In contrast YAP1, perhaps on a different temporal scale, sets a different regulatory potential for response to incoming signals, including availability of TEAD transcription factors^53^ that can interact with the actin polymerization responsive protein VGGL4 which rises in response to cytochalasin D^50^. In addition, in mesenchymal stem cells, nuclear deformation itself can promote YAP1 nuclear localization, confirming that nuclear mechanics will affect cell function^54^. As such, the type of mechanical input – dynamic versus static – and the force induced effect on nuclear structure may pre-set either a β-catenin or YAP configuration to prime subsequent cellular responses to incoming signals.

In contrast to β-catenin, YAP1 exits the nucleus as actin flows inward due to cytochalasin D induced cytoplasmic F-actin depolymerization^18^. Stable cytoplasmic F-actin polymerization in response to a stiff substrate, on the other hand, leads to YAP1 nuclear translocation where F-actin forces on the nuclear surface cause flattening of the nucleus and opening of nuclear pores to support YAP1 transfer^14^. Here, LPA-induction of cytoplasmic F-actin affected YAP1 but not β-catenin shuttling. Here our study showed that while dynamic strain increased both total cell and nuclear stiffness more than did static strain (^~^2-fold) it did not result in YAP1 nuclear entry. Dynamic low intensity vibration (LIV), in comparison, promoted YAP1 nuclear entry only when the cell stiffness increased by 4-fold^16, 55^. These contrasting relationships to mechanical input highlight that β-catenin and YAP1 represent principal components of two functionally distinct and non-redundant pathways by which cytoskeletal mechanics modulate nuclear functions linked to proliferation and differentiation of stem cells.

Nuclear import for β-catenin, YAP1, or β-actin occurs in the absence of classic nuclear localization signals (NLS)^24, 25^. As such, all three proteins depend on other factors for targeted nuclear import, including association with structural elements and co-transporters. For example, disabling LINC complex function impedes mechanically-induced nuclear import of β-catenin under dynamic strain^17^ and YAP1 under static strain^56^. Further, while YAP1 force-induced nuclear entry is independent of GTPase, inhibiting GTPases has been shown to decrease its nuclear accumulation^14^. TAZ which represents a YAP1 co-partner, does appear to have a non-canonical NLS which may affect YAP1 transfer^25^. Other studies have shown that YAP1 can be rapidly imported via interactions with transcription factors (e.g., Runx2) that exhibit highly dynamic nuclear/cytoplasmic compartmentalization^57, 58^. Nuclear import of actin requires cofilin/importin-9 interaction with monomeric and dimeric actin^40^. Because cofilin-1 does contain a NLS^47^, this protein enables actin nuclear transport in concert with importin-9; knockdown of either cofilin-1 or importin-9 blocks actin transport^18^. The latter finding was corroborated here by visualization of YFP-labeled actin. Here we present the novel finding that knockdown of either cofilin-1 or importin-9 also prevents nuclear transfer of β-catenin. This supports that nuclear import of β-catenin depends on strain-dependent interactions with mono- and dimeric actin shuttled into the nucleus through cofilin/importin-9 dependent mechanisms.

## Conclusion

It is remarkable that two key molecules involved in controlling cell state decisions, β-catenin and YAP1, respond so specifically to different types of physical forces. A key question that emerges is how their presence in the nucleus controls cell function. In the case of mesenchymal stem cells, neither protein is linked to a clearly defined intracellular signaling pathway. We propose that β-catenin, through its association with actin dynamics, results in a structural orchestration of gene responses to the constant environmental barrage of incoming signals.

---

**Supplemental 1.**
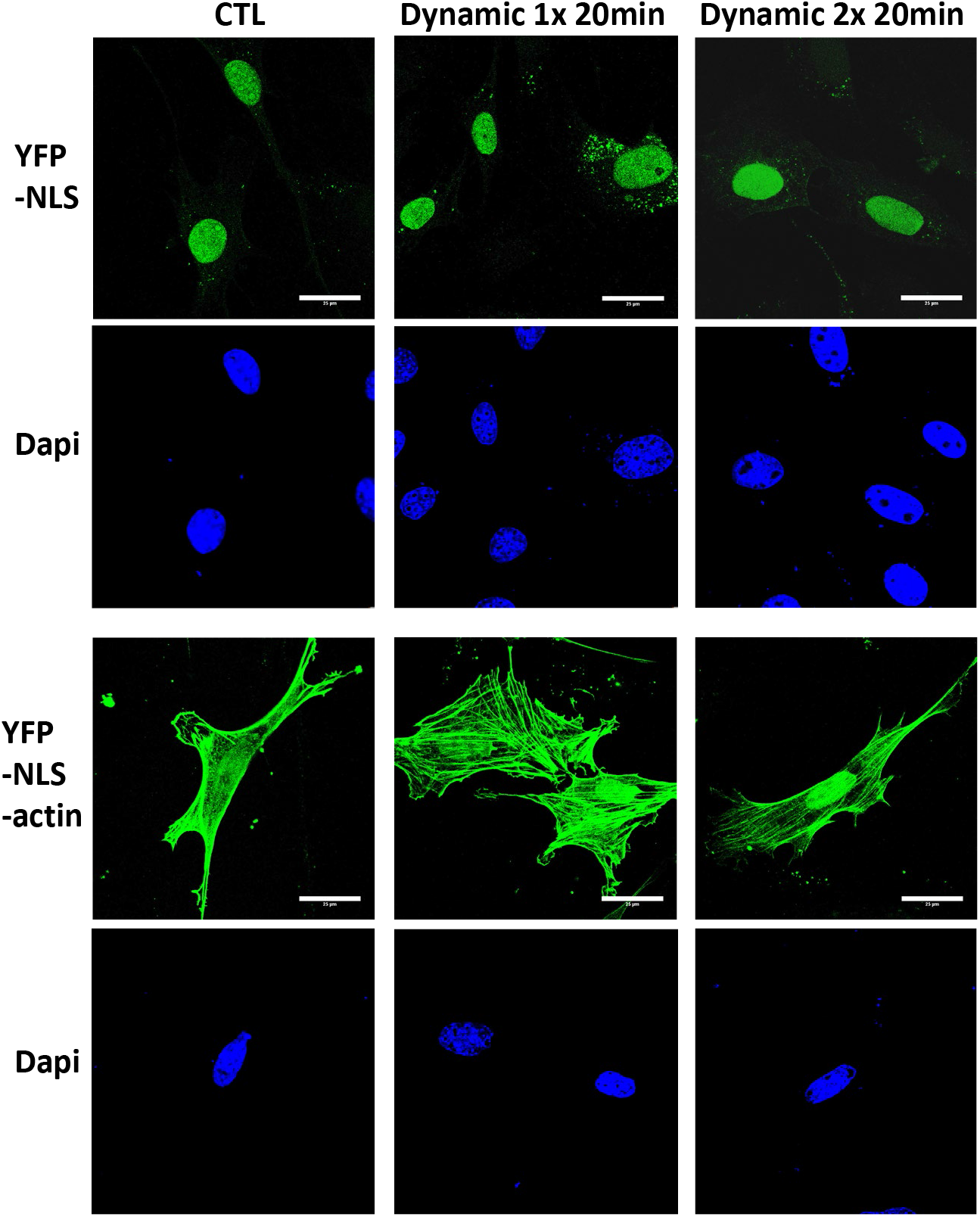
YFP-NLS is intranuclear while YFP-NLS-Actin is found in the cytoplasm. YFP-NLS (Addgene, Cat#145274) and YFP-NLS-βActin (Addgene, Cat#60613) were overexpressed in MSC and applied single or double time of 2% dynamic strain same as that in fig 3, scale bar = 25 μm. More YFP-NLS-actin is seen intranuclear after the second bout of strain.

**Supplemental 2.**
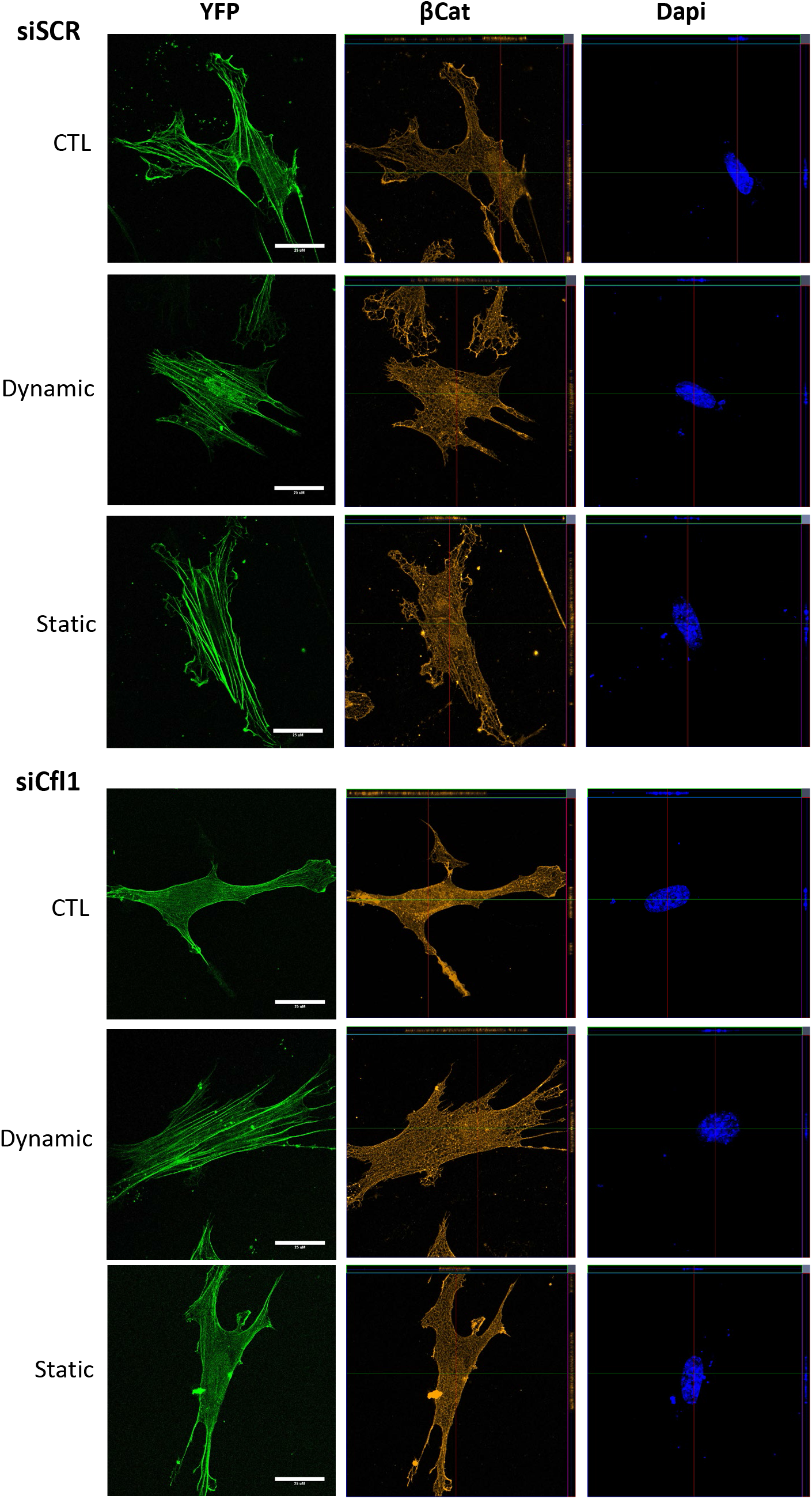
Dynamic vs static strain in YFP-expressing cells). MSC overexpressing YFP-NLS-βActin (Addgene, Cat#60613) in +/- knock-down of *Cfl1* or *Ipo9* were subject to 2% dynamic or % static strain before stain with anti-β-catenin (yellow) and DAPI (blue), scale bar = 25 μm.

## Abbreviations

RT-qPCR: reverse transcriptase real time quantitative polymerase chain reaction
GFP: green fluorescent protein
YFP: yellow fluorescent protein

## Data availability

The data underlying this article are available in the article and in its online supplementary material.

---

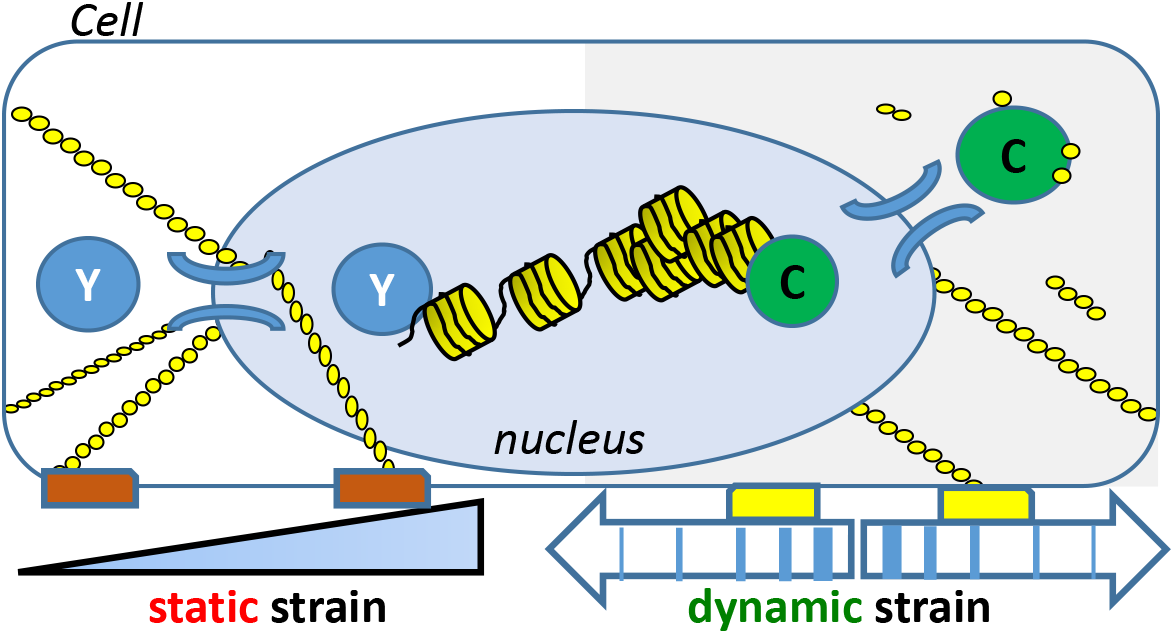
β-Catenin (C) entry into the nucleus due to dynamic strain is dependent on actin transport. YAP (Y) nuclear entry depends on a static actin-LINC structure delivering force to open nuclear pores.

